# Computationally-guided tuning of ligand sensitivity in a GPCR-based sensor

**DOI:** 10.1101/2021.09.21.461282

**Authors:** Daniel Keri, Reto B. Cola, Zacharoula Kagiampaki, Patriarchi Tommaso, Patrick Barth

## Abstract

Genetically-encoded fluorescent sensors for neuromodulators are increasingly used molecular tools in neuroscience. However, these protein-based biosensors are often limited by the sensitivity of the protein scaffold towards endogenous ligands. Here, we explored the possibility of applying computational design approaches for enhancing sensor sensitivity. Using the dopamine sensor dLight1 as proof of concept, we designed two variants that boost the sensor’s potency (EC50) for dopamine and norepinephrine by up to 5- and 15-fold, respectively. Interestingly, the largest effects were obtained through improved designed allosteric transmission in the transmembrane region of the sensor. Our approach should prove generally useful for enhancing sensing capabilities of a large variety of neuromodulator sensors.

## Main Text

Investigating the spatiotemporal dynamics of neuromodulator release with high resolution has been historically limited by several factors. A range of tools has been established in the past, that provided either high temporal resolution (e.g. fast-scan cyclic voltammetry), high spatial resolution (e.g. pHluorins), or high molecular specificity (e.g. microdialysis). Only recently, the development of GPCR-based genetically-encoded fluorescent sensors drastically expanded the technological landscape by allowing in vivo neuromodulator imaging with high spatiotemporal resolution and molecular specificity^1^. Since more than 400 GPCRs are known and the general design strategy of such GPCR-based sensors has proven versatile^2^, several new molecular tools have been developed to study the release properties of dopamine, norepinephrine, acetylcholine, serotonin, ATP, endocannabinoids and adenosine, among others ^3^. While the number of such GPCR-based sensors has been increasing at a rapid pace, modalities for tuning sensor potency for their endogenous ligands, which largely determines the in vivo sensitivity and selectivity of the probes, have remained largely unexplored. Improving receptor’s sensitivity and response to ligand binding has proven difficult through traditional site-directed mutagenesis because possible design solutions often involve non intuitive long-range mutational effects that cannot be easily inferred from protein structures. To address this limitation, computational approaches have been developed recently to predict and design sequence variations that modulate the allosteric signal properties of GPCRs^4^. Here, we combined this computational strategy with traditional protein engineering approaches to design and validate sensor variants with improved ligand sensitivity.

For this proof-of-principle study, we opted to improve the ligand sensitivity of the dopamine sensor dLight1.3b. This sensor is the variant with largest dynamic range from the dLight1 suite of dopamine sensors^2^ with a max ΔF/F_max_ of 930%, but has a relatively low affinity (EC_50_ above 1 μM). It was engineered by replacing the third intracellular loop (ICL3) of the dopamine receptor D1 with a circularly permutated green fluorescent protein (cpGFP), followed by mutagenesis and high-throughput of the amino acid linker regions flanking the fluorescent protein. By focusing our computational efforts onto redesigning the orthosteric sites or allosteric regions, that have previously not been targeted by traditional site-directed mutagenesis, we herewith introduce a novel approach for altering potencies of GPCR-based sensors in a previously unexplored manner (**Fig. 1A**).

**Figure 1:**
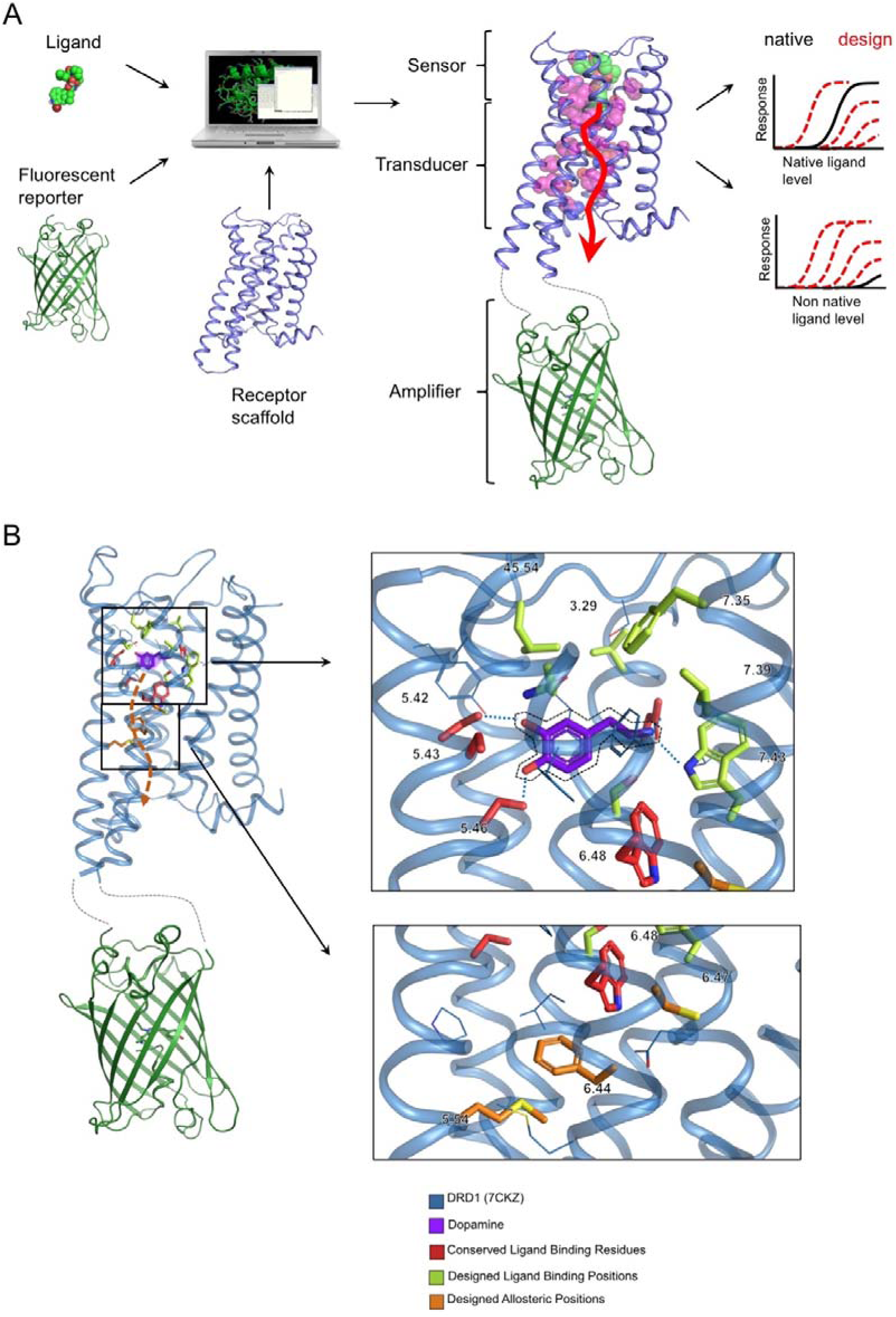
Computational design of GPCR-based fluorescent sensors with enhanced sensitivity. **A**. Schematic description of the computational design approach. Biosensors are built using 3 distinct molecules: the ligand to detect; the GPCR sensing the ligand and transmitting the signal to the reporter; and the fluorescent amplifier. Ligand orthosteric binding site and allosteric transmission regions are designed to improve the sensor response to ligand binding. **B**. Cartoon representation of dopamine bound dLight1.3b highlighting the positions targeted for design. Residue positions are indicated using the Ballesteros Weinstein notation.

The study was performed before the dopamine D1 structures were released. The inactive and active state structures of dLight1.3b bound to dopamine were modeled by homology to those of dopamine D2 and B2AR using the method IPHoLD^5^ (**Methods**). We identified the residues most frequently found in contact with dopamine in our D1 models. These included conserved residues on TMH3 and 5 identified by mutagenesis to be important for ligand binding as well as more variable residues on ECL2, TMHs 3, 6 and 7 that we targeted for design (**Fig. 1B**). By analogy to the dopamine D2 receptor, we identified also putative allosteric sites that couple the extracellular to the intracellular regions and control the responses of the receptor to ligand binding. These sites include microswitches in the core regions of TMHs 5 and 6 that undergo conformational changes upon ligand binding and were also targeted by our computational approach (**Fig. 1B**). Variants in the orthosteric binding site predicted to increase binding affinity to dopamine were selected using the Rosetta Ligand design approach^6^ and those predicted to enhance allosteric response were selected using our allosteric design pipeline^4^.

Mutations with predicted enhanced sensitivity for dopamine were then cloned into the dopamine sensor dLight1.3b backbone, expressed in HEK cells and characterized *in vitro* for fluorescent sensor responses. None of the designed mutations in the orthosteric site significantly improved sensor responses and we often observed reduced sensitivity compared to the original dLight1.3b. By contrast, close to 50% of our allosteric designs displayed increased dopamine sensitivity (**Fig. 2A**). Detailed ligand dose titration of the fluorescence responses showed that the potency increased by up to 5- and 15-fold for dopamine and norepinephrine, respectively (**Fig. 2B**). These large sensitivity enhancements are remarkable considering that dLight1.3b is based on the D1 receptor that is already naturally evolved to sense both dopamine and norepinephrine. Both D1 designs F6.44M and C6.47L target the highly conserved PIF and CWxP motifs, respectively, which are typically considered to be optimal for allosteric signaling. Consistent with our previous observations on the dopamine D2 receptor, the results suggest that the ligand sensitivity of naturally evolved GPCRs are far from optimal and can readily be improved through the rational design of allosteric mutations at highly variable as well as conserved regions.

**Figure 2:**
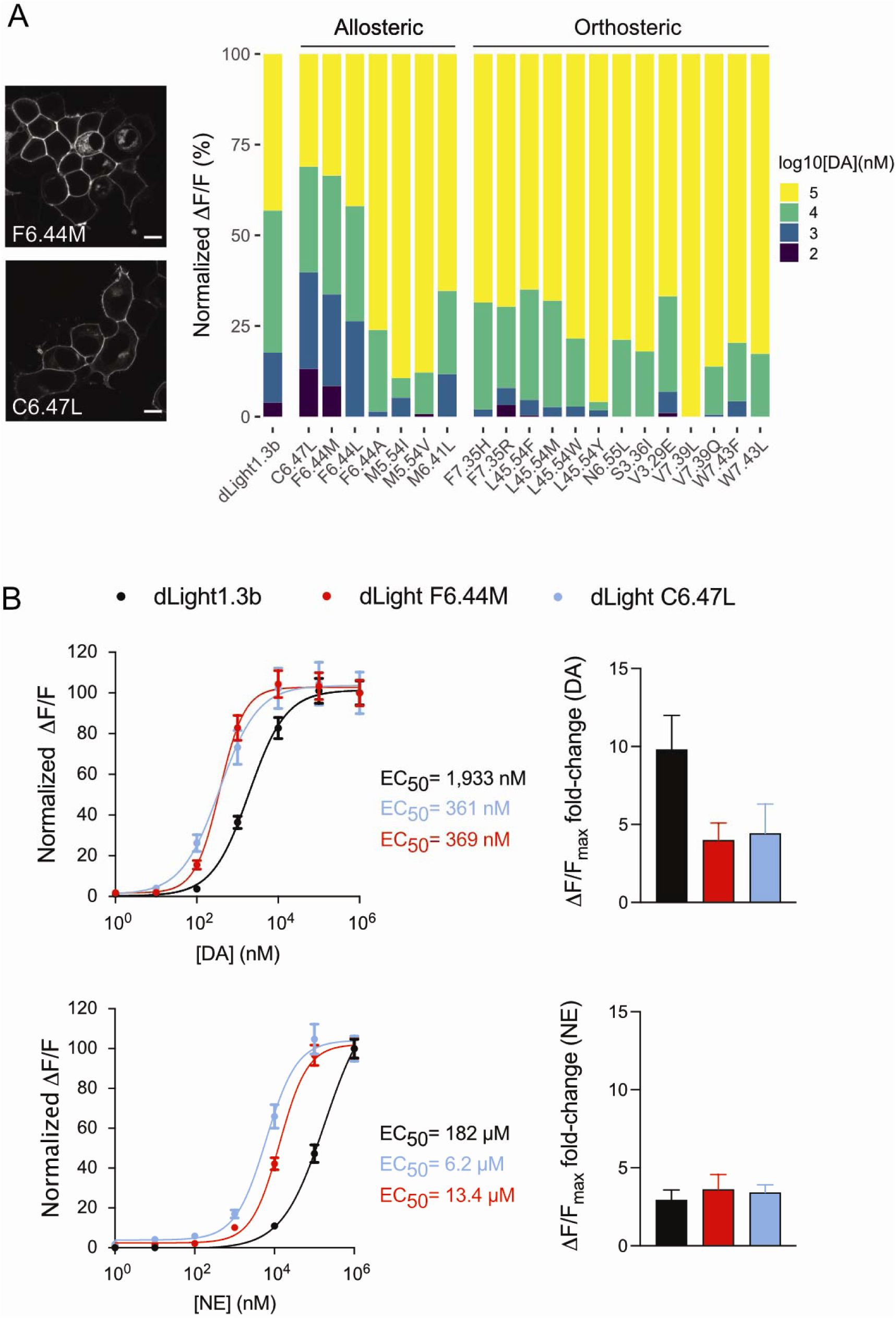
*In vitro* characterization of designed dopamine sensors. **A**. An initial screen with 4 different DA concentrations per construct was performed to identify candidates with increased normalized ΔF/F response to low concentrations. Only mutations with positive responses were considered for this screen. The two mutations C6.47L and F6.44M in allosteric positions, both showed increased normalized ΔF/F responses compared with dLight1.3b already at 100nM [DA]. None of the modelled mutations in orthosteric positions resulted in improved responses to low [DA]. Left, representative images of sensor expressing HEK cells are shown on the left. Scale bar, 10 μm. **B**. The two identified mutants from the initial screen in A) were further analyzed in more detail. The overlaid dose-response curves for the two mutants and dLight1.3b confirms the improvement in potency for both DA and NE. While the absolute dynamic range (max ΔF/F_max_) was unaltered between constructs in the response to NE, there was a marked reduction in absolute dynamic range for the two high-affinity mutants. Amino acid positions are referred to by the Ballesteros-Weinstein numbering convention of the D1 receptor. [DA] = dopamine concentration; [NE] = norepinephrine concentration.

Interestingly, the non-conserved allosteric microswitch residues 5.54 and 6.41, involved in a key TMH5 and TMH6 contact, highlight a site-specific differential optimization of allosteric signal transmission between the D1 and D2 receptors. Mutagenesis of the D1 native M5.54, M6.41 towards the native D2 T5.54, L6.41 by way of the D1 M6.41L design results in a loss in sensitivity (ΔLog(EC_50_) = 0.98), whereas the previously blindly predicted D2 T5.54M design results in a gain in sensitivity (ΔLog(EC_50_) = -1.20)^7^; suggesting a higher local optimality for the native D1 sequence. Despite such differences in local optimality, all dopamine (including D1 and D2) receptor sensitivities are surprisingly very similar^2,8-10^. These observations suggest that there is a pressure of selection for maintaining globally suboptimal signaling properties. One potential contributing factor to this may be the need for high signaling selectivity. Consistent with this hypothesis, the enhanced sensitivity observed in designed receptors was obtained at the expense of their selectivity. The sensitivity of designed D1s to norepinephrine reached ∼90% of that for dopamine versus 30% for WT (**Fig. 2B)** and corroborate similar observations with D2 designs and the weak partial agonist serotonin. Hence, native dopamine receptors may have favored the minimization of noise (i.e. high selectivity) from structurally similar agonists at the expense of sensitivity. Whether allosteric signaling and selectivity can be reprogrammed independently remains a promising future prospect to be investigated.

While the allosteric designs represent a clear success of our approach, we also sought to understand the origin of the disagreement between the orthosteric binding site predictions and the experimental outcome. We analyzed the dopamine D1 models and compared them to the recently solved cryoEM structure. With up to 90% native contact recovery, our models match the experimental structure very closely and are unlikely at the origin of the low design success rate. The computational ligand binding design method that we used was extensively validated in various systems including soluble proteins^6^ and GPCRs^5^ but often involved the design of new packing or polar interactions with the ligands. Creating such contacts with dopamine or norepinephrine onto the D1 scaffold was challenging because both ligands are very small and already engaged through polar interactions with the native receptor. Hence, solutions for improving binding through direct receptor-ligand contacts were scarce and are reminiscent of failed previous attempts at improving small ligand binding to B2AR^11^.

Overall, our study suggests that our computational allosteric design approach can be applied to reprogram the sensitivity of GPCR-based sensors through designed allosteric pathways. Since the *in silico* calculations are very efficient (1 to 3 CPU hours per site), the approach should prove useful for reducing the effort in screening novel variants with fine-tuned responses and allow more rapid expansion of the toolbox of receptor-based ligand sensors for cutting edge *in vivo* detection applications. Lastly, our method provides a rational blueprint for systematically exploring and better understanding the evolution of allosteric properties in GPCRs.

## Methods

### Computational modeling

Homology models of the dopamine D1 receptor active state were created from the recently solved dopamine D2 receptor Gi-bound structure (6VMS), and the epinephrine-bound B2 adrenergic receptor structure (4LDO). The D1 inactive state homology model was based on the D2 inactive structure (6CM4). The generation of all models was based on previously established protocols^4,5^. Sequential steps of coarse-grained docking, structure relaxation, relaxed model clustering and final high resolution docking were carried out using iPHoLD^2^. 10,000 ligand-bound decoys were generated using coarse-grained ligand docking, loop modeling and structure relaxation. The top 10% lowest scoring models were clustered and the cluster center from the 10 largest clusters used as starting scaffolds for high resolution ligand docking simulations. For each starting cluster center, 10,000 ligand-bound models were generated and the top 10% lowest scoring decoys were selected for ligand binding mode clustering via a DBSCAN-based algorithm using ligand heavy atom coordinates. For each cluster center model, the lowest energy model from the largest ligand cluster was used for design.

### Computational design

Allosteric designs were generated by performing in silico deep scanning mutagenesis on the aforementioned homology models at predetermined allosteric hub sites using established protocols^4,12^. Designs were selected based on each mutation’s ability to maximize the difference in allosteric dynamic coupling between the active and inactive state models while minimizing changes in both active and inactive state stabilities.

Orthosteric ligand binding designs were generated using Rosetta’s Ligand design protocol^6^. Residues within 8Å of the docked ligand were designated as design positions, with the exception of the highly conserved TM3 aspartate, TM5 serines, and the TM6 toggle switch tryptophan. We performed in silico deep scanning mutagenesis at the design and adjacent positions to discover both single or combination affinity-enhancing mutations. Typically 100 decoys were generated to ensure score convergence. Designs with improved ligand binding interface energy and no significant destabilization of the receptor structure relative to WT were selected. Models meeting these criteria were ran through the ligand docking procedure again to ensure the designed mutations do not promote alternate ligand binding modes and were filtered out otherwise.

### Molecular cloning

Cloning of dLight1.3b and related mutant constructs was performed on the background of a *pCMV_dLight1*.*1 (Addgene #111053)* expression vector plasmid. Mutations were cloned into the expression vector by polymerase chain reaction (PCR) using a *PFUultra II Hotstart PCR Master Mix* (Agilent Technologies). The obtained PCR products were subsequently treated with *DPNI* (New England Biolabs) and cleaned up with the *Qiaquick PCR Purification Kit* (Qiagen). The blunt-ended linear PCR products were then phosphorylated with *T4* polynucleotide kinase (ThermoFisher Scientific) and ligated overnight with *T4 ligase* (New England Biolabs). The ligated product was then transformed into E.coli 10beta (New England Biolabs) and colonies were grown on LB-agar plate containing kanamycin for selection. Individual colonies were picked and inoculated overnight in kanamycin-containing 2YT medium. DNA extraction from bacterial clones was performed with the *Qiaprep Spin Miniprep Kit* (Qiagen) and DNA sequences were verified with Sanger sequencing (Microsynth).

### Cell culture and confocal imaging

HEK293T cells (ATCC CRL-3216) were grown and maintained in culture media containing Dulbeco’s Modified Eagle Medium (DMEM, Gibco), 10% fetal bovine serum (FBS, Gibco), and 1x antibiotic-antimycotic (Gibco) at 37°C and supplemented with 5% CO_2_. For passaging cells and plating into 35mm glass-bottom dishes (MatTek), Trypsin 0.05% EDTA (Gibco) was added to the cells and blocked with equal volume of FBS-containing medium after detachment. Suspended cells were then centrifuged for 3 minutes at 150x G and the supernatant was removed. The cell pellet was re-suspended in 10mL of fresh culture media (37°C) and cells density was calculated with a TC20 automated cell counter (BioRad). 100’000 to 150’000 cells were plated per 35mm glass-bottom dish and transfected the following day. Transfection was performed with the *Effectene Transfection Reagent* (Qiagen) according to the supplier’s manual and a media change was performed 6h after transfection. Two days after transfection, the cells were imaged at a confocal microscope (Zeiss LSM 800) and washed 1x with *Hank’s Balanced Salt Solution* (Gibco). Dopamine titrations were performed by directly application of dopamine solutions to the dish at the indicated concentration.

### Data analysis

Confocal imaging was analyzed within ImageJ. In short, the time lapse images were thresholded for pixel intensity and ROIs of cell membranes were selected. The mean pixel intensities for each ROI was then extracted for each frame of the time lapse and saved as a .csv file. Calculation of ΔF/F_max_ for each ROI in each frame was performed with custom written Matlab codes. Plotting of dose-response curves was subsequently performed in Prism Graphpad, while data analysis for data obtained in the initial screening procedure was performed in R.

## Acknowledgments

D.K. and P.B. were supported by an SNF grant (number 31003A_182263), funding from EPFL and the Ludwig Institute for Cancer Research. T.P. was supported by funding from the European Research Council (ERC) under the European Union’s Horizon 2020 research and innovation program (Grant agreement No.s 891959 and 101016787), as well as by an SNF grant (number 310030_196455).

